# Lobule-specific dosage considerations for cerebellar transcranial direct current stimulation during healthy aging – a computational modeling study using age-specific MRI templates

**DOI:** 10.1101/535658

**Authors:** Zeynab Rezaee, Anirban Dutta

**Affiliations:** Department of Biomedical Engineering, University at Buffalo, Buffalo, USA

## Abstract

The world population aged 60 years and older is expected to double between 2015 and 2050. Aging is associated with a decline in cognitive and motor performances which are a part of geriatric syndromes. Aging is also associated with the loss of cerebellar volume where the cerebellum has a considerable contribution in cognitive and motor functions. Therefore, cerebellar transcranial direct current stimulation (ctDCS) has been proposed to study and facilitate cerebellar function during aging. However, the one-size-fits-all approach used for ctDCS can lead to variability in the cerebellar lobule-specific dosing due to age-related changes in the cerebellar structure. Therefore, we investigated lobular electric field (EF) distribution during healthy aging for age groups of 18 to 89 years where computational modeling was based on age-appropriate human brain magnetic resonance imaging (MRI) templates (http://jerlab.psych.sc.edu/NeurodevelopmentalMRIDatabase/). A fully automated open-source pipeline (Realistic vOlumetric-Approach to Simulate Transcranial Electric Stimulation – ROAST) was used for the age-group specific EF modeling. Then, we extracted the EF distribution at the 28 cerebellar lobules based on a spatially unbiased atlas (SUIT) for the cerebellum. Our computational results showed that the EF strength increased significantly at certain important cerebellar lobules (e.g., Crus I and Crus II relevant for cognitive function) contralateral (contra) to the targeted (ipsi) cerebellar hemisphere at an older age that reduced the ctDCS specificity. Specifically, two-way ANOVA showed that the lobules as well as the age-group (and their interaction term) had a significant effect (p<0.01). Post-hoc multiple comparison tests at Alpha=0.01 using Bonferroni critical values showed that Right (Ipsi) Crus I, Right (Ipsi) Crus II, Right (Ipsi) VI, Vermis VIIb, Vermis VIIIa, Right (Ipsi) VIIb, Left (Contra) VIIIb, Left (Contra) IX, Right (Ipsi) VIIIa, Right (Ipsi) VIIIb, Vermis VIIIb, Right (Ipsi) IX, and Vermis IX, and the age-group 18, 18.5, 19, 20-24, 45-49, 50-54, 70-74, 75-79, 85-89 years experienced higher electric field strength (>0.11V/m). Since there is a dichotomy between the sensorimotor cerebellum and the cognitive cerebellum, therefore, subject-specific MRI based head modeling for lobule-specific dosage considerations will be necessary for clinical translation of ctDCS to address geriatric cerebellar syndromes.

## Introduction

The world population aged 60 years and older is expected to increase to 2 billion by 2050 from 900 million in 2015 (Ageing and health). Older age leads to a decline in cognitive and motor performances which are a part of geriatric syndromes. Non-invasive brain stimulation (NIBS) technique using the small magnitude of the direct electrical current, called transcranial direct current stimulation (tDCS), has been proposed for combating age-related cognitive decline (Woods, 2017). Ongoing clinical study on “augmenting cognitive training in older adults (The ACT Study)” proposes tDCS of the frontal cortices using the one-size-fits-all approach with electrodes placed at F3 and F4 (10-20 EEG system) (Woods et al., 2018). Since frontal lobe atrophy is implicated in patterns of age-related cognitive decline (Hogan et al., 2011) therefore tDCS of the frontal cortices is warranted. Nevertheless, recent literature suggests that cerebellar morphology may be better at predicting cognitive and motor performances (Bernard and Seidler, 2014) with a high concentration of neurons that contributes to both motor and cognitive processes. Such age-related volume loss in the cerebellum is associated with a decline in the motor and cognitive functions, such as working memory, processing speed, and spatial processing (Koppelmans et al., 2017). Therefore, cerebellar tDCS (ctDCS) (Grimaldi et al., 2016) is proposed as a novel approach for understanding cerebellar function during aging where studies using ctDCS showed various benefits in both motor and cognitive behaviors (see a review of (Ferrucci and Priori, 2014)). Galea and colleagues (Galea et al., 2009) delivered ctDCS using two sponge electrodes (5cmx5cm) with one electrode centered at 3 cm lateral to the inion and the other electrode positioned on the right buccinator muscle. They showed a polarity-specific modulation of cerebellar brain inhibition (CBI) with a current intensity of 2 mA applied for 25 minutes where CBI is a marker of neuroplasticity. Here, one-size-fits-all ctDCS approach (called “Celnik montage” henceforth (Galea et al., 2009)) is postulated to be inadequate in ensuring comparable lobule-specific dosing across subjects due to inter-subject variability of the complex cerebellar structure with distinct motor and non-motor regions related to distinct motor, language, social, working memory, and emotional functions (van Dun et al., 2018). This is crucial since cerebellar cortex is organized into a myriad of functional units called microzones with segregated loops between the cerebellum and prefrontal cortex, parietal cortex, paralimbic cortex and superior temporal sulcus (Lawrenson et al., 2018). Table 1 lists the cerebellar region, the associated area in the brain, and the behavioral task from literature.

**Table 1:**
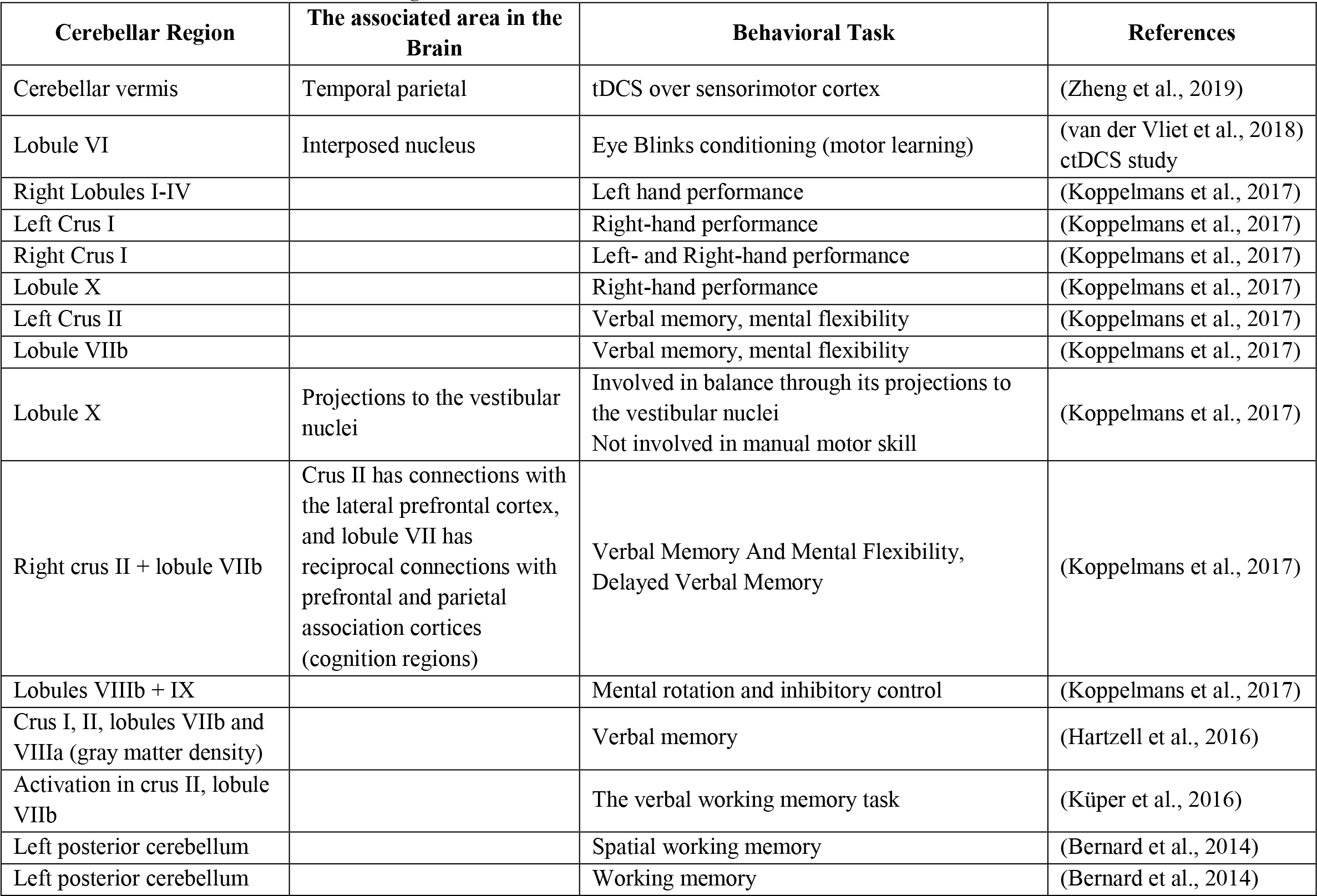
Cerebellar region, associated areas in the brain, and the behavioral task

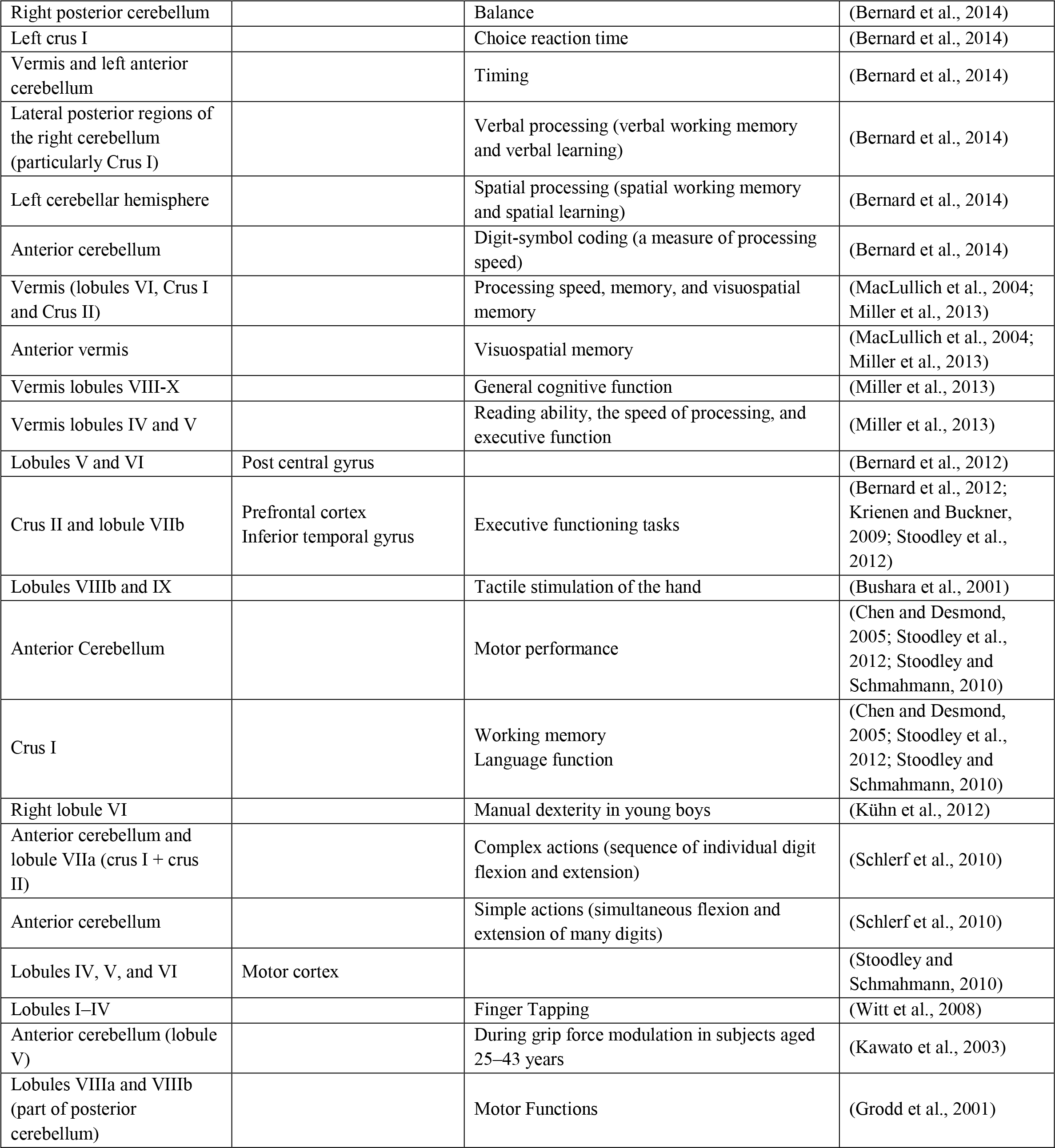

Modeling studies (Rampersad et al., 2014) have shown that the ctDCS delivered with “Celnik montage” leads to a fairly unilateral electrical field over the anterior and posterior cerebellar hemisphere in the 25-29 years age range in most human ctDCS studies (Ferrucci et al., 2008, 2013; Ferrucci and Priori, 2014; Galea et al., 2011; Sparing et al., 2009). However, due to the changes of cerebellar morphology even in healthy aging (Bernard and Seidler, 2014), we postulate that one-size-fits-all approach for ctDCS developed for a young adult brain would lead to a significant difference in the lobular electric field distribution in older adults than young adults. Specifically, aging leads to a reduction in cerebellar volume and white matter integrity where some cerebellar lobules are more prone to shrink than others during aging. For example, both posterior and anterior vermis display pronounced age-related volume loss (Raz N et al., 1992; Shah et al., 1991), as does the anterior lobe (Bernard and Seidler, 2013; Escalona et al., 1991; Luft et al., 1999; Shah et al., 1991). Other studies found that the most impacted region is the superior posterior cerebellum (i.e., lobules Crus I, Crus II, VI, and VIIb) (Abe et al., 2008; LEE et al., 2005; Paul et al., 2009). Hence, there is significant inter-subject variability in the impact of age on the shrinkage of the cerebellar lobules that may be associated with the decline in motor and cognitive behavior. There is a dichotomy between the sensorimotor cerebellum and the lateral portions of the cerebellum (cognitive cerebellum) (Lawrenson et al., 2018) that needs to be accounted for in optimizing ctDCS dose to address the decline in motor and cognitive behavior during aging. For example, lobules IV, V, VI, and VIII were shown related to motor functions while lobules VI and VII were involved in non-motor functions (van Dun et al., 2018). Here, lobule-specific dosage considerations for ctDCS in older adults can be based on computational modeling of electrical fields, e.g., one performed in children that advised caution (Kessler et al., 2013). However, a lobule-specific electrical field distribution was not presented by the computational modeling study by Rampersad and colleagues (Rampersad et al., 2014) that is postulated to be crucial for ctDCS due to the dichotomy between the sensorimotor cerebellum and the cognitive cerebellum (Lawrenson et al., 2018). Moreover, since cerebellar cortex has a very high concentration of neurons with highly organized distribution, therefore, computation of lobule-specific electric field distribution can be very effective for ctDCS dosage considerations to address different cerebellar syndromes (Lawrenson et al., 2018), e.g., the vestibulocerebellar syndrome, the cerebellar motor syndrome, and the cerebellar cognitive affective syndrome (Koziol et al., 2014).

In this computational modeling study, we postulate that the investigation of the changes in the electrical field distribution across cerebellar lobules during healthy aging will help us to better understand the lobule-specific dosage considerations of ctDCS in older adults. Therefore, we investigated the lobular electric field distribution across eighteen age groups from 18 to 89 years of age by applying our published computational pipeline (Rezaee and Dutta, 2019) on age-appropriate human brain magnetic resonance imaging (MRI) templates (http://jerlab.psych.sc.edu/NeurodevelopmentalMRIDatabase/). Our computational modeling study leveraged Realistic volumetric Approach to Simulate Transcranial Electric Stimulation (ROAST) (Huang et al., 2018) for head modeling and finite element analysis (FEA), and then applied Spatially Unbiased Infratentorial Template for the Cerebellum (SUIT) atlas (Diedrichsen et al., 2009) to divide the cerebellum into 28 lobules. In this paper, we focused our analysis on the cerebellar regions which were reported to be more affected during the aging process, e.g., the vermis area of the anterior part of the cerebellum (mostly related to sensorimotor) and Crus I and Crus II (for cognitive performance) (Koziol et al., 2014; Stoodley et al., 2012). To our knowledge, there is no systematic investigation of the lobular electric field distribution during ctDCS in an aging population. Since the effects of brain atrophy on the current density distribution in the brain have been shown recently (Mahdavi and Towhidkhah, 2018) so new methodologies are necessary for dosing ctDCS for the older adults. This is also important for safety considerations since ctDCS is being increasingly applied on older adults (Woods et al., 2018), and up to 4mA current intensity has been presented (Chhatbar et al., 2018). In principal accordance, we present a rational approach for lobule-specific dosage considerations for ctDCS and investigate the ctDCS dosing with “Celnik montage” during healthy aging.

## Material and Methods

### Computational Modeling Pipeline: Head Model Creation

To investigate age-related effects on the electric field distribution of the human cerebellar lobules, the age-appropriate human brain MRI templates were obtained online at http://jerlab.psych.sc.edu/NeurodevelopmentalMRIDatabase/ with the permission of Dr. John Richards. The data consisted of average T1-weighted MRI for the head and brain, and segmenting priors for gray matter (GM), white matter (WM), and cerebrospinal fluid (CSF) for the eighteen groups of averaged adult heads (male and female), ranging from 18-89 years of age, from the Neurodevelopmental MRI Database (Escalona et al., 1993; Fillmore et al., 2015; Richards et al., 2016; Richards and Xie, 2015; Sanchez et al., 2012). For the first four age-groups, the age intervals were six months (i.e., 18-18.5-19-19.5 years of age), and then from age 20 to 89 years for the next fourteen age-groups, the average T1-weighted MRI were grouped by an interval of 5 years. We applied our computational pipeline (Rezaee and Dutta, 2019) on the average T1-weighted MRI data that leveraged Realistic volumetric Approach to Simulate Transcranial Electric Stimulation (ROAST) (Huang et al., 2018) – a Matlab script based on three open source software; Statistical Parametric Mapping (SPM) (Penny et al., 2011), Iso2mesh (Fang and Boas, 2009), and getDP (Dular et al., 1998). Here, average T1-weighted MRI for each age group was used to construct the age-group specific head models for finite element analysis (FEA). ROAST used SPM8 (SPM - Statistical Parametric Mapping) to segment the head and brain. After segmentation, five tissues were achieved for the tetrahedral volume mesh, namely, Scalp, Skull, Cerebrospinal Fluid (CSF), Gray Matter (GM), and White Matter (WM). These different brain tissues for the volume mesh were modeled as different volume conductors for MEA in the ROAST. Here, isotropic conductivity used for the different brain tissues (Windhoff et al., 2013) were (in S/m): Scalp=0.465; Skull=0.01; CSF=1.654; GM=0.276; WM=0.126 (Datta et al., 2009; Foerster et al., 2018; Windhoff et al., 2013).

### Finite element analysis of ctDCS using Celnik montage and age-group specific head model

The creation of the tetrahedral volume meshing of the head model and the FEA was performed using ROAST pipeline (Huang et al., 2018), which provided a powerful numerical tool to solve the required partial differential equations (PDE), as shown in Figure 1. FEA divides the head model (volume conductor) into spatial elements and nodes (tetrahedral volume mesh) for discrete computations of the PDE (electric field). Here, ctDCS was modeled based on “Celnik montage,” where we applied 2mA current intensity using two sponge electrodes (5cm×5cm) with the 5cm × 5cm anode placed over the right cerebellum, 1 cm below and 3 cm lateral to the inion (Iz, 10/10 EEG system)(Galea et al., 2009), and the 5cm × 5cm cathode placed over the right buccinator muscle. The electrodes were modeled in ROAST as saline-soaked sponges placed at the given scalp locations using 10/10 EEG system (Giacometti et al., 2014). FEA was conducted on each age-group specific head model to compute the ctDCS induced electric field in the brain tissues. In all the simulations, the voxel size was 1mm^3^.

**Figure 1.**
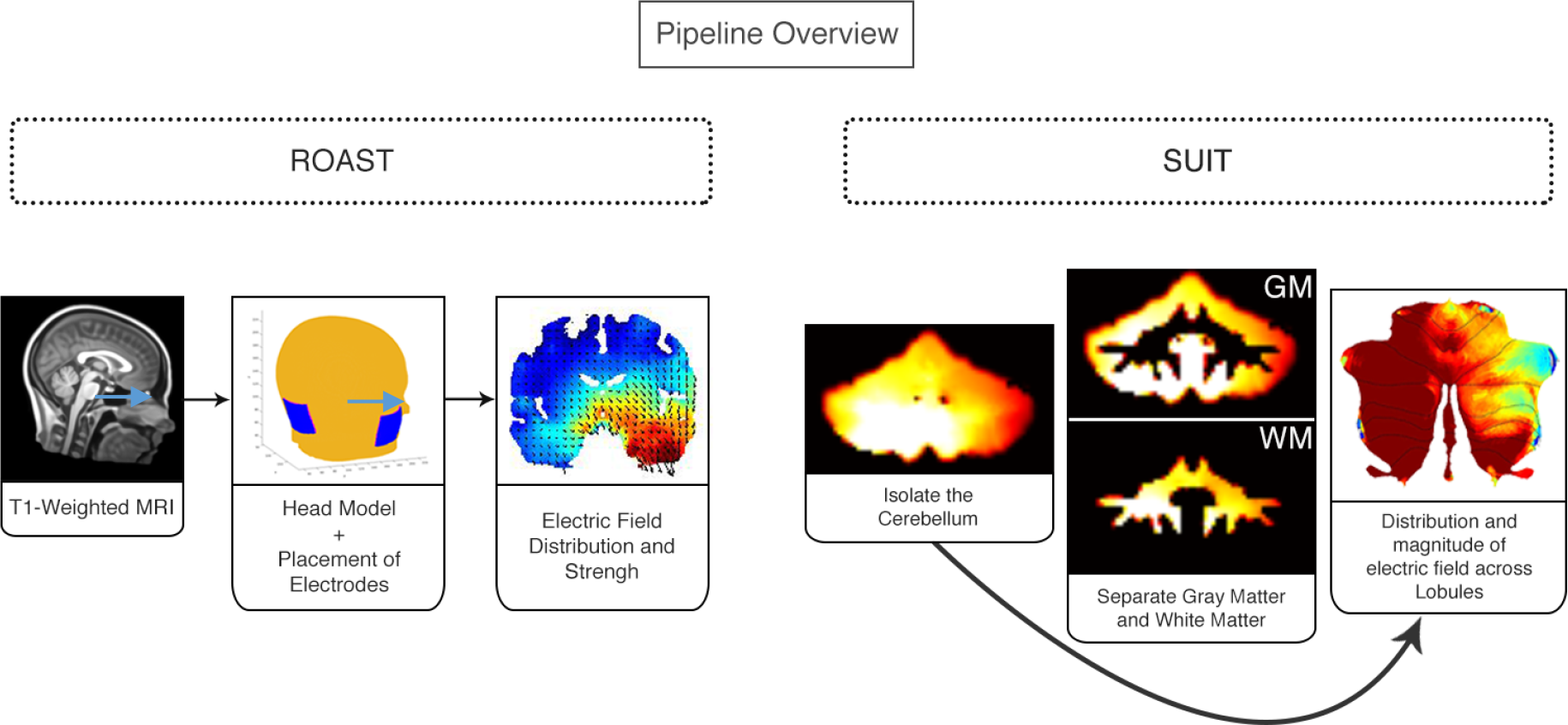
Overview of our computational pipeline. The workflow starts with the average MRI of each age group obtained online at http://jerlab.psych.sc.edu/NeurodevelopmentalMRIDatabase/ with the permission of Dr. John Richards. ROAST toolbox (Huang et al., 2018) was used to create head models for the 18 available age groups. ROAST places the electrodes in the given position on the scalp and performs finite element analysis (FEA). Celnik montage with the 5cm × 5cm anode placed over the right cerebellum, 1 cm below and 3 cm lateral to the inion (Iz, 10/10 EEG system), and the 5cmx5cm cathode placed over the right buccinator muscle was used for ctDCS with ±2mA current intensity. The isolation of the cerebellum was performed using SUIT toolbox (Diedrichsen et al., 2009). The cerebellum was isolated including only gray matter (GM). Lobular electric field distribution was visualized in a 2D map (called flatmap).

### Isolation of the cerebellar lobules and flatmap visualization of the electric field distribution

To investigate the electric field distribution in the cerebellar lobules, we first isolated the cerebellar electric field distribution using the age-group specific cerebellar mask computed by ROAST. The recent version of ROAST provides the electric field distribution as NIFTI (Neuroimaging Informatics Technology Initiative) images. Then, the electric field distribution in the gray matter (GM) and white matter (WM) were extracted from the NIFTI images. Then, the cerebellar lobules were isolated using SUIT pipeline in SPM (http://www.fil.ion.ucl.ac.uk/spm/). T1 images for each age-group were reoriented into LPI (neurological) orientation. We used the cerebellar lobules defined in the SUIT atlas (Diedrichsen, 2006; Diedrichsen et al., 2009), which consisted of the left and right cerebellar hemispheres and the vermis. The isolation map was manually verified in an image viewer by the researcher. After the isolation, the cerebellum was normalized to the SUIT atlas template using the cropped image and the isolation map. A nonlinear deformation map to the SUIT template is the result of the normalization step. The image was resampled into the averaged age-group space. The volume of the cerebellar lobules, defined by the SUIT atlas (Diedrichsen, 2006), was used for the extraction of the lobular electric field distribution. As part of the SUIT toolbox, a flat representation of the human cerebellum is provided to visualize data after volume-based normalization. Therefore, flatmap script in the SUIT toolbox was used to visualize the GM electric field distribution in the cerebellar lobules, as shown in Figure 1.

Rampersad and colleagues (Rampersad et al., 2014) have shown a fairly unilateral electrical field over the anterior and posterior cerebellar hemisphere during ctDCS based on “Celnik montage,” however did not provide the lobule-specific electrical field (EF) distribution. Therefore, we investigated the hemispheric specificity during healthy ageing for the related cerebellar lobules (Left I_IV, Right I_IV; Left V, Right V; Left VI, Right VI; Left Crus I, Right Crus I; Left Crus II, Right Crus II; Left VIIb, Right VIIb; Left VIIIa, Right VIIIa; Left VIIIb, Right VIIIb; Left IX, Right IX; Left X, Right X) across the eighteen age-group. Specifically, we investigated the ratio of the EF (in V/m) distribution in targeted (ipsi) and contralateral (contra) cerebellar hemisphere to capture the specificity:

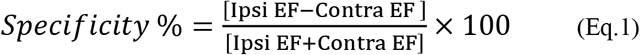

Here, the mean value of the EF at each lobule was used for calculating this ratio.

### Statistical analysis of the electric field distribution across eighteen age groups ranging from 18-89 years

The electric field was computed at all the voxels of the 28 cerebellar lobules defined by the SUIT atlas (Diedrichsen, 2006). Then, we analyzed the GM electric field distribution across eighteen age-group specific head models using two-way ANOVA (‘anovan’ in Matlab) for the factors of interest – lobules, age-group, and their interactions. In the Generalized Linear Model (GLM), the proportion of the total variability in the dependent variable that is accounted for by the variation in the independent variable was found using the eta-squared effect size measure. Post-hoc multiple comparison tests were conducted using Bonferroni critical values.

## Results

The SUIT flatmap for the cerebellar lobules is shown in Figure 2A and the ROAST head model for the “Celnik montage” is shown in Figure 2B. The ROAST head model (Figure 2B) was used to compute the SUIT flatmap (Figure 2A) for the electric field distribution in the cerebellar gray matter, as shown in Figure 2C. The lobular electric field strength across the age-group (shown as flatmap in Figure 2C) had a maximum value of 0.3V/m.

**Figure 2.**
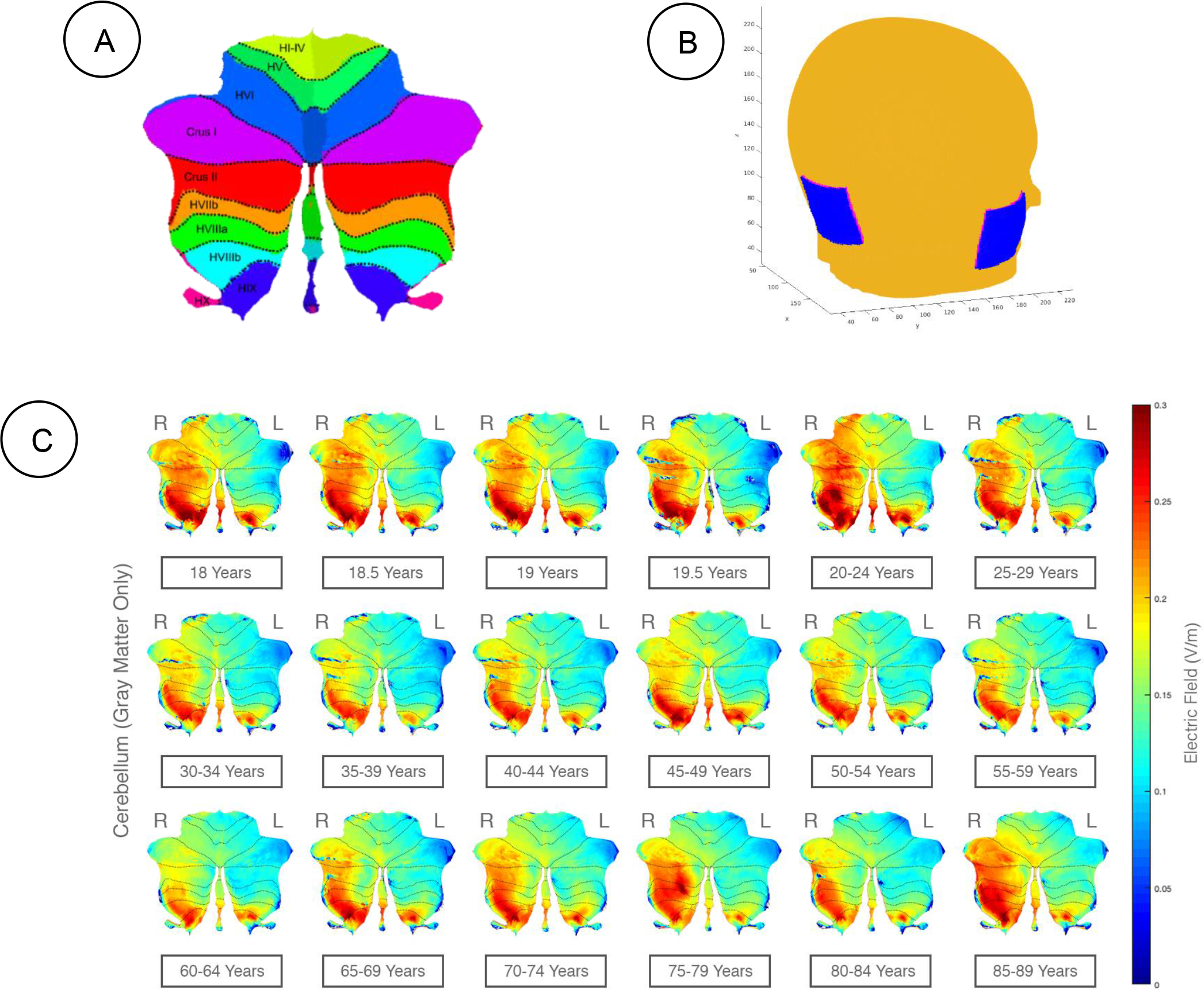
Visualization of electric field strength (V/m) of ctDCS in different age groups in flatmap (SUIT toolbox) **A) Cerebellar Lobules.** Cerebellar lobules showed using a 2D representation (called flatmap) based on SUIT atlas (Diedrichsen, 2006). **B) Head model in ROAST.** The figure is a representation of the head model, electrode placement for Celnik montage, and the (x,y,z)direction which were used for all age groups. **C) The electric field strength in the cerebellar gray matter where right (R) hemisphere is the targeted side (ipsi) and the left (L) hemisphere is the contralateral side (contra).** ctDCS using Celnik Montage with a direct current of 2mA was simulated using our computational pipeline, and the lobular electric field (magnitude) distribution across age-groups is visualized on a 2D flatmap. Right (R) and Left (L) cerebellar hemispheres are specified in the figure. **D) The electric field strength in the contralateral (left) lobules (e.g., Crus I, Crus II, and VIIb) increased with aging.**

Appendix 1 presents the lobular electric field distribution (as 2D flatmap) in the X, Y, Z directions in the cerebellar gray matter and Appendix 2 presents the GM electric field strength (V/m) across all the cerebellar lobules. Here, two-way ANOVA showed that the lobules, age-group as well as their interaction term had a significant effect (p<0.01) on the GM electric field distribution (ANOVA table shown in Figure 3A). Post-hoc multiple comparison tests at Alpha=0.01 were conducted using Bonferroni critical values which showed that Right (Ipsi) Crus I, Right (Ipsi) Crus II, Right (Ipsi) VI, Vermis VIIb, Vermis VIIIa, Right (Ipsi) VIIb, Left (Contra) VIIIb, Left (Contra) IX, Right (Ipsi) VIIIa, Right (Ipsi) VIIIb, Vermis VIIIb, Right (Ipsi) IX, and Vermis IX experienced higher electric field strength (>0.11V/m - lower limit in precision (Huang et al., 2017)) across all the age-group and age-group 18, 18.5, 19, 20-24, 45-49, 50-54, 70-74, 75-79, 85-89 years experienced the higher electric field strength (>0.11V/m - lower limit in precision (Huang et al., 2017)) across all the lobules, as shown in Figure 3B and 3C respectively.

**Figure 3.**
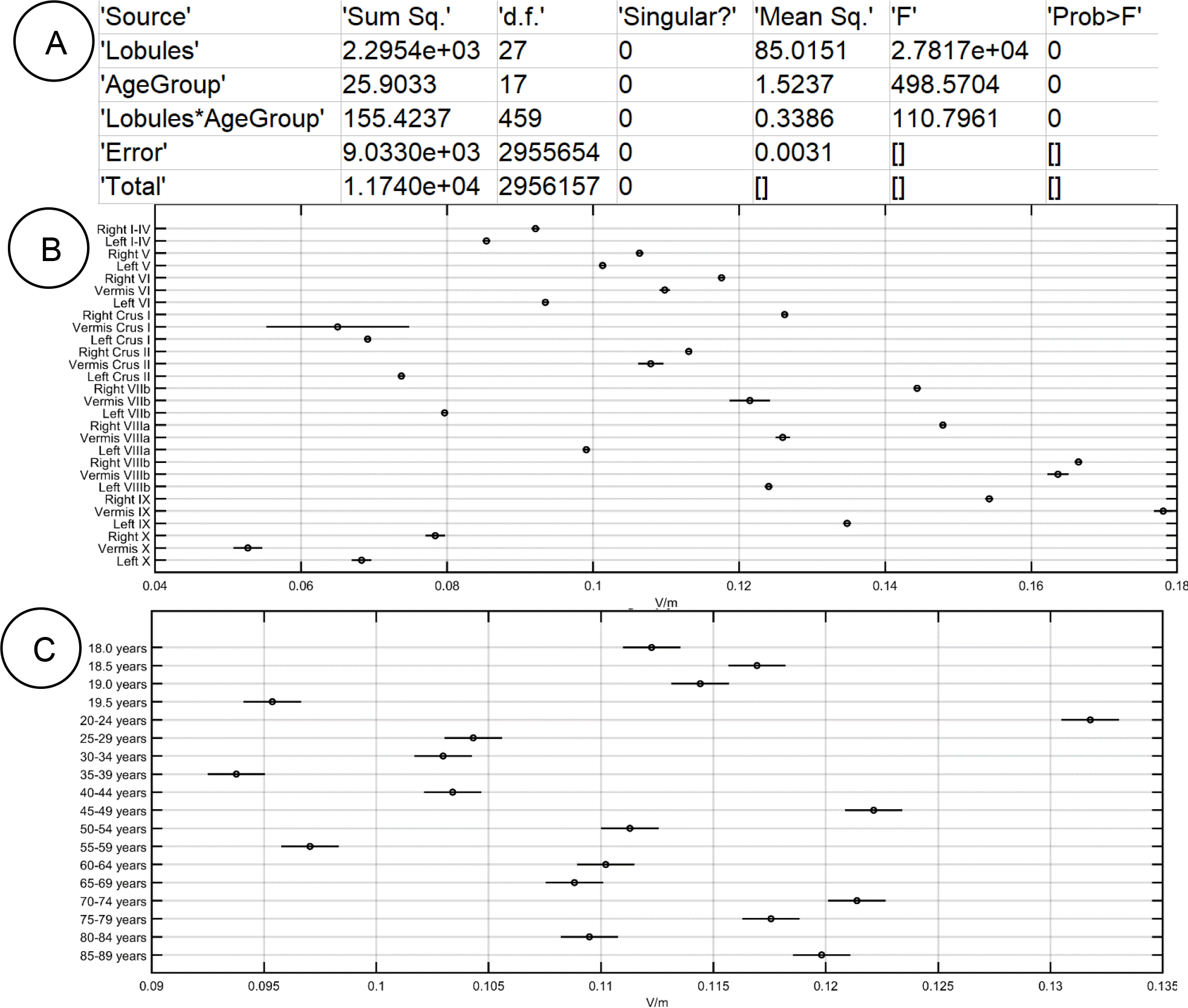
Statistical Results. Post-hoc multiple comparison tests at Alpha=0.01 were conducted using Bonferroni critical values where right (R) hemisphere is the targeted side (ipsi) and the left (L) hemisphere is the contralateral side (contra) **A) ANOVA table.** The p-value 0 indicates that the mean responses across levels for the factors ‘Lobules’ and ‘AgeGroup’, as well as their interaction term represented by ‘Lobules*AgeGroup’, are significantly different. **B) Across AgeGroup.** Right Crus I, Vermis VIIb, Vermis VIIIa, Right VIIb, Left VIIIb, Left IX, Right VIIIa, Right VIIIb, Vermis VIIIb, Right IX, and Vermis IX experienced highest electric field strength (>0.12V/m). **C) Across Lobules.** Age-group 20-24, 45-49, 70-74 years experienced the highest electric field strength (>0.12V/m).

It was found that the GM electric field strength (or magnitude) in some of the contralateral (left) lobules increased with aging (see Appendix 2). Specifically, Crus I, Crus II, and VIIb showed a significant increase as shown in Figure 4A, that can lower the specificity of the ctDCS. Indeed, the most aging impacted region is the superior posterior cerebellum (i.e., lobules Crus I, Crus II, VI, and VIIb) (Abe et al., 2008; LEE et al., 2005; Paul et al., 2009), therefore, the specificity in the GM electric field strength in those lobules (i.e., lobules Crus I, Crus II, VI, and VIIb) were investigated across the eighteen age-group, as shown in Figure 4B. Also, the Table 2 presents the specificity values for all the relevant cerebellar lobules (Ipsi I_IV, Contra I_IV; Ipsi V, Contra V; Ipsi VI, Contra VI; Ipsi Crus I, Contra Crus I; Ipsi Crus II, Contra Crus II; Ipsi VIIb, Contra VIIb; Ipsi VIIIa, Contra VIIIa; Ipsi VIIIb, Contra VIIIb; Ipsi IX, Contra IX; Ipsi X, Contra X) across eighteen age groups. Table 2 shows that the specificity decreased with aging in certain lobules (e.g., lobules Crus I, Crus II, and VIIb) while improved in other lobules (e.g., I_IV, and X).

**Table 2:**
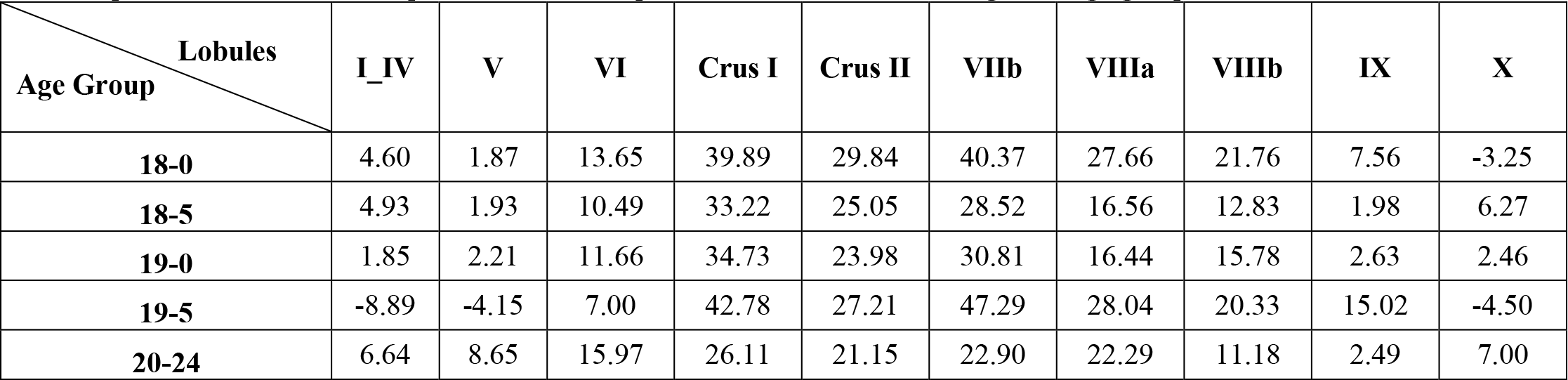
Specificity % (eqn. 1) values for all the relevant lobules (Ipsi I_IV, Contra I_IV; Ipsi V, Contra V; Ipsi VI, Contra VI; Ipsi Crus I, Contra Crus I; Ipsi Crus II, Contra Crus II; Ipsi VIIb, Contra VIIb; Ipsi VIIIa, Contra VIIIa; Ipsi VIIIb, Contra VIIIb; Ipsi IX, Contra IX; Ipsi X, Contra X) across the eighteen age groups.

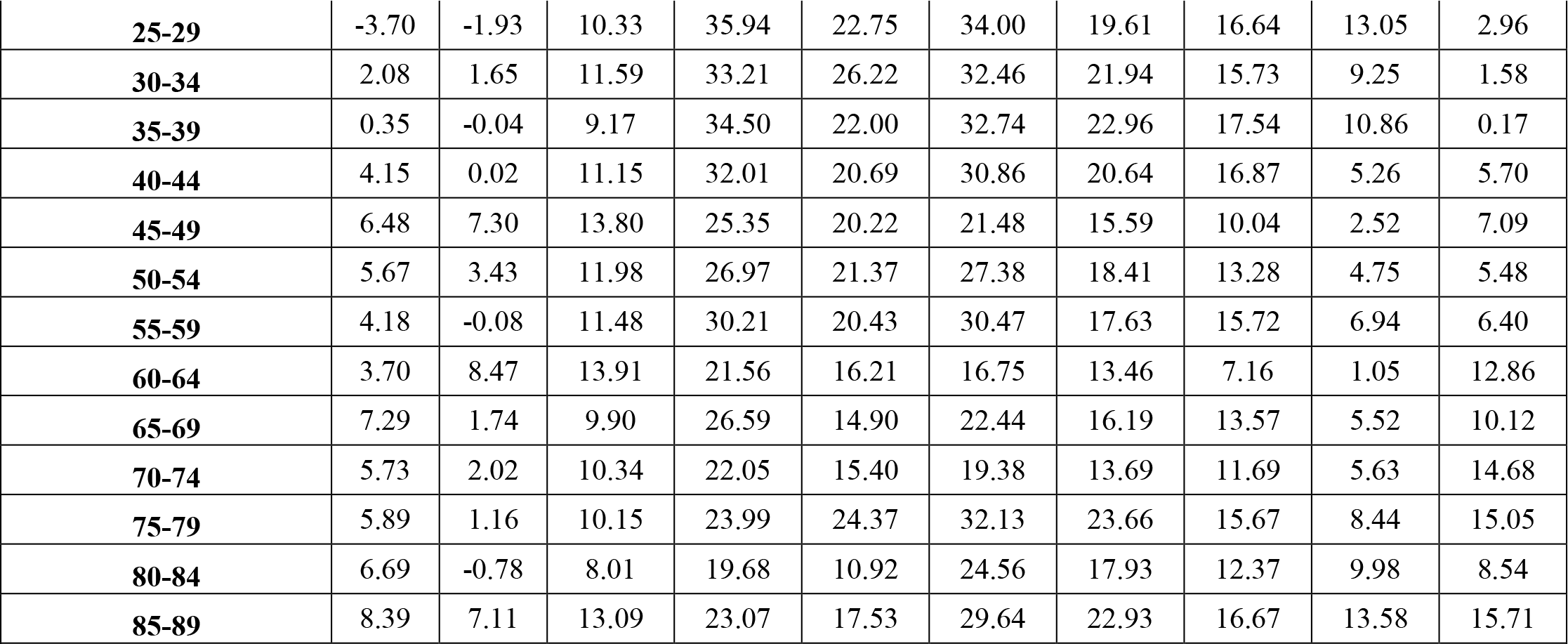

**Figure 4.**
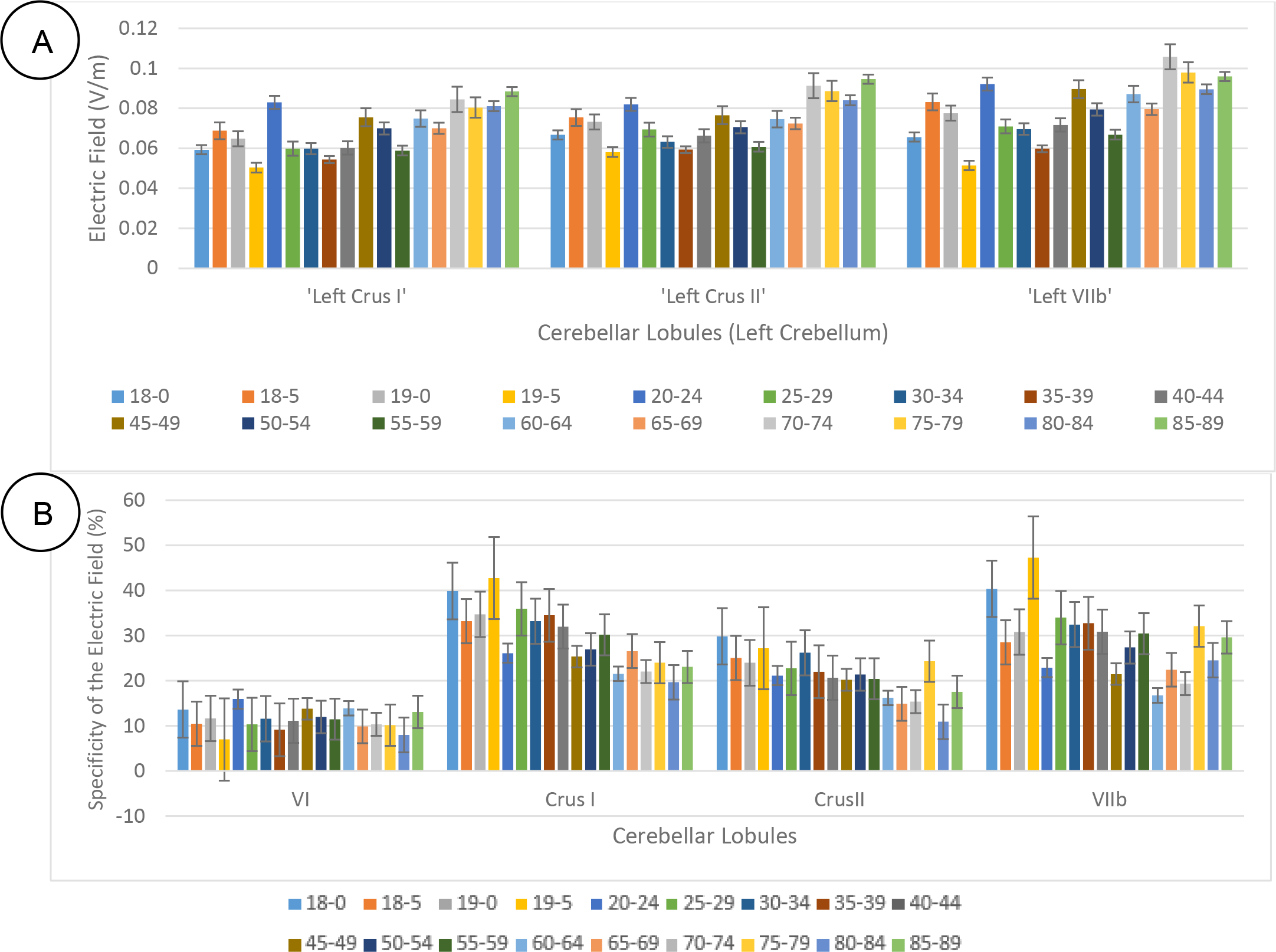
A) The electric field strength in some of the contralateral (left) lobules increased with aging. B) Electric field specificity (eqn. 1) for the cerebellar lobules – Crus I, Crus II, VI, and VIIb – across the eighteen age-group (shown with color). Electric field magnitude in the cerebellar lobules Crus I, Crus II, VI, and VIIb in the contralateral cerebellar hemisphere increased with aging, i.e., the specificity decreased with aging. The plot bar shows the Specificity for the specific lobules (Crus I, Crus II, VI, and VIIb). The standard error is also shown on the bar graph.

## Discussion

The cerebellum is one of the most affected regions of the brain by aging (Raz and Rodrigue, 2006) and is a better predictor of performance decline in older adults than the prefrontal cortex (Bernard and Seidler, 2014). Therefore, the relationship between the volume of cerebellar regions with behavior functioning has been used to understand the neurobiology of age-related changes in motor and cognitive performances (Koppelmans et al., 2017). The main goal of this study was to investigate the lobular electric field distribution applied by the ctDCS using “Celnik montage” in age-specific head models ranging from 18-89 years of age. We performed a comparison among the eighteen age groups to study the effect of aging on the ctDCS electric field distribution across the cerebellar lobules. Although the electric field distribution for ctDCS using Celnik montage remained mostly unilateral (Figure 2C), however, there was a decrease in specificity with aging (Figure 4). Moreover, the electric field strength in some cerebellar lobules contralateral to ctDCS targeted hemisphere, e.g., contra Crus I, contra Crus II, and contra VIIb (see Figure 4A), showed an increase in younger (18, 18.5, 19, 20-24 years of age) and older (45-49, 50-54, 70-74, 75-79, 85-89 years of age) adults that also experienced higher electric field strength (>0.11V/m - lower limit in precision (Huang et al., 2017)) across all lobules (see Figure 3C). This result supports the prior findings that showed brain volume increased in young adulthood, then the increase continued slowly and reached a plateau in the 4th decade, and then decreased in the old age (Bartzokis et al., 2001; Jernigan et al., 2001; Raz et al., 2005). This shrinkage in cerebellar cortex and white matter volume starts from around the 40s (Walhovd et al., 2011). In our study, the increase in the electric field strength in the contralateral (non-targeted) cerebellar hemisphere can be explained by current shunting due to an increase in CSF thickness and a decline in gray matter volume with aging that has been discussed earlier (Mahdavi and Towhidkhah, 2018). Here, cortical thinning and GM atrophy will have a net effect of increasing the distance between the cortical surface and the scalp. This decline in GM volume has been shown by the volumetric results which showed anterior and superior posterior part of the vermis to be more susceptible in the aging process (Bernard and Seidler, 2013). As CSF thickness increases, there is more current dispersion to the contralateral (non-targeted) cerebellar hemisphere in older adults. Although Celnik montage has been shown to have unilateral effects (Rampersad et al., 2014), however, our results showed that aging leads to spill-over of electric field outside of the targeted brain region. The electric field strength across the eighteen age groups showed that the spill-over of the electric field increased mostly after 70 years of age (Figure 4A). Here, even though “Celnik montage” primarily targeted the posterior and the inferior parts of the right cerebellum, as shown by the SUIT flat map results in Figure 2C, however, the specificity decreased with aging in certain important lobules of cognitive cerebellum as shown in Figure 4B and Table 2.

The age-related volume decline is more evident in anterior cerebellar lobe (lobule I-IV and lobule V) and cerebellar superior posterior lobe (Crus I, Crus II, lobule VI, and lobule VIIb) compared to the inferior posterior lobe (lobule VIIIa, VIIIb, and IX) (Bernard and Seidler, 2013; Hulst et al., 2015; Koppelmans et al., 2017). Therefore, evidence-based application of ctDCS needs careful lobule-specific dosage considerations in older adults when most healthy human studies are conducted in the 25-29 years age range (Ferrucci et al., 2008, 2013; Ferrucci and Priori, 2014; Galea et al., 2011; Sparing et al., 2009). A recent computational modeling tDCS study on aging (Mahdavi and Towhidkhah, 2018) investigated the current density distribution of tDCS with brain atrophy using two montages for the motor area. In both the montages, the simulation results for the younger head model was more focal under the electrode and across brain folds. Indeed, as the GM volume declines with age, the CSF thickness between the scalp to gray matter increases. This increase in the CSF thickness leads to more current dispersion in aging. Here, an inverse square law relationship between electromagnetic field strength and the distance. Although the tDCS technique has not shown serious adverse effects (Ferrucci et al., 2015) and the benefits of tDCS has been shown in rehabilitation of older adults (Block and Celnik, 2012; Ferrucci and Priori, 2014; Galea et al., 2011; Raz et al., 2013; Reis et al., 2008), the computational modeling study (Mahdavi and Towhidkhah, 2018) discussed possible harmful effects in the older adults brain in the absence of optimization. Specifically, the current density in the deeper brain regions increased in, the older adult brain that may be susceptible to overstimulation due to tDCS protocols designed for younger adults (Mahdavi and Towhidkhah, 2018). Here, a challenge specific to dosage considerations for ctDCS during ageing is due to the non-linear decline in the total cerebellar volume where the volume loss seems to continue faster in advanced age and by a quadratic function (Hoogendam et al., 2012; Luft et al., 1999; Walhovd et al., 2005, 2011). Moreover, vermis and anterior lobe volume seem to decline with a logarithmic pattern while in the posterior lobe a quadratic function explained the volume loss (Bernard et al., 2014). Such heterogeneous volume loss across cerebellar lobules will shunt the ctDCS current in a complex manner where subject-specific neuroimaging based FEA is necessary for dosage considerations. Furthermore, GM and white matter (WM) volume change with a different pattern during aging (Ge et al., 2002) where there is a linear decrease in GM percentage with age whereas there is a quadratic relation between WM volume and age. Here, WM deterioration during aging is an important part of the cerebellum volume loss in advanced age (Andersen et al., 2003) (Sullivan et al., 2010). Also, intracranial space for both GM and WM were found to be significantly less in older adults that can also lead to more electric field dispersion (Good et al., 2001; Lemaître et al., 2005).

The electric field dispersion outside of the targeted brain region during aging (Mahdavi and Towhidkhah, 2018) require dosage considerations in older adults. Here, an age-specific head model for FEA is needed to optimize ctDCS in the older adults due age-related cerebellar volume loss to address different cerebellar syndromes (Lawrenson et al., 2018), e.g., the vestibulocerebellar syndrome, the cerebellar motor syndrome, and the cerebellar cognitive affective syndrome (Koziol et al., 2014). For example, the motor performance is associated with the volume of the anterior lobe and top part of the superior posterior lobe, and the cognitive performance is related to the volume of the bottom part of the superior posterior lobe and the inferior posterior lobe (Koppelmans et al., 2017). Moreover, most recent works show that lobules IV, V, VI, and a part of VIII are related to motor functions (van Dun et al., 2018) while lobules VI, VII, VIIIa, Crus I and Crus II (Hartzell et al., 2016; Koppelmans et al., 2017; Küper et al., 2016; Stoodley et al., 2012; van Dun et al., 2018) are involved in non-motor functions, therefore, lobule-specific electric field modeling is necessary in the older adults to avoid spill-over effects on non-targeted functions. Here, a positive correlation between cognitive performance and vermis size was found in older adults (MacLullich et al., 2004; Miller et al., 2013). Also, larger volumes resulted in better performance (Koppelmans et al., 2017; Nyberg et al., 2012). Furthermore, cerebellar cortex is organized into a myriad of functional units, called microzones, that will require optimization of ctDCS induced electric field direction for the modulation of synchronous activity within and between cerebellar micro-complexes (De Gruijl et al., 2014). Such detailed modeling is crucial due to the complex structure of the cerebellum and also because a small change within 0.05-0.5V/m can induce reproducible and noticeable changes in neuron activity (Bogaard et al., 2019).

There are some limitations in our study that should be taken into account. Investigating the effects of aging on the brain current flow needs accurate brain tissue factors, e.g., age-related reductions in skull conductivity (Hoekema et al., 2003), which was considered constant during aging in this study. Another limitation of this study is the use of averaged age-appropriate human brain magnetic resonance imaging (MRI) templates (http://jerlab.psych.sc.edu/NeurodevelopmentalMRIDatabase/) for the construction of age-specific head models. Here, the gender was not considered separately for the computational head modeling. Although, there are studies which reported age-related brain changes to vary with gender (Cowell et al., 1994; Ge et al., 2002; Xu et al., 2000) and the male and female cerebellar lobe are different in size (Liu et al., 2016), no gender-specific correlation between cerebellar volume have been reported (Arani et al., 2015; Jäncke et al., 2015). Another limitation of this study is that our MRI age-group data had five years interval. However, cerebellar volume decrease has been reported in shorter intervals (Raz et al., 2005; Smith et al., 2007). In future studies, subject-specific computational analysis of lobular electric field distribution followed by neurophysiological testing of the aging effects on ctDCS based on the cerebellum brain inhibition (CBI) (Fernandez et al., 2018) can confirm our approach. Here, CBI is currently explained by the Purkinje cell-mediated inhibition of facilitation of the motor cortex (M1). Therefore, early ctDCS of the cerebellar GM can activate the Purkinje cell, which inhibits the deep cerebellar nuclei (DCN)-thalamo-cortical WM pathway (van Dun et al., 2018) and may prevent the WM loss in advanced age (Ge et al., 2002; Jernigan et al., 2001). Indeed, WM volume loss in both cerebrum and cerebellum starts later than GM volume loss (Bernard et al., 2014; Jernigan et al., 2001). Therefore, ctDCS targeting the Right (Ipsi) Crus I, Vermis VIIb, Vermis VIIIa, Right (Ipsi) VIIb, Left (Contra) VIIIb, Left (Contra) IX, Right (Ipsi) VIIIa, Right (Ipsi) VIIIb, Vermis VIIIb, Right (Ipsi) IX, and Vermis IX with high electric field strength (>0.12V/m) across age-group may prevent the decline in older adults in certain cerebellar lobules (Vermis VI, Crus I, and X) that have a strong association with cerebellar WM volume decline.

## Author Contributions

ZR conducted the computational modeling under the guidance of AD. All the authors have drafted the work and revised it critically and have approved the final version before submission.

## Acknowledgment

ZR and AD would like to acknowledge the fruitful discussions with German collaborators, Dr. Giorgi Batsikadze and Dr. Dagmar Timmann, Department of Neurology, Essen University Hospital, University of Duisburg-Essen, Essen, Germany, and Prof. Dr. med. Michael Nitsche, head of the department of psychology and neurosciences, Leibniz Research Centre for Working Environment and Human Factors, Germany.

## Funding

University at Buffalo SUNY funded this computational work.

## Conflict of Interest Statement

The authors declare that they have no conflict of interest.

**Appendix 1.**
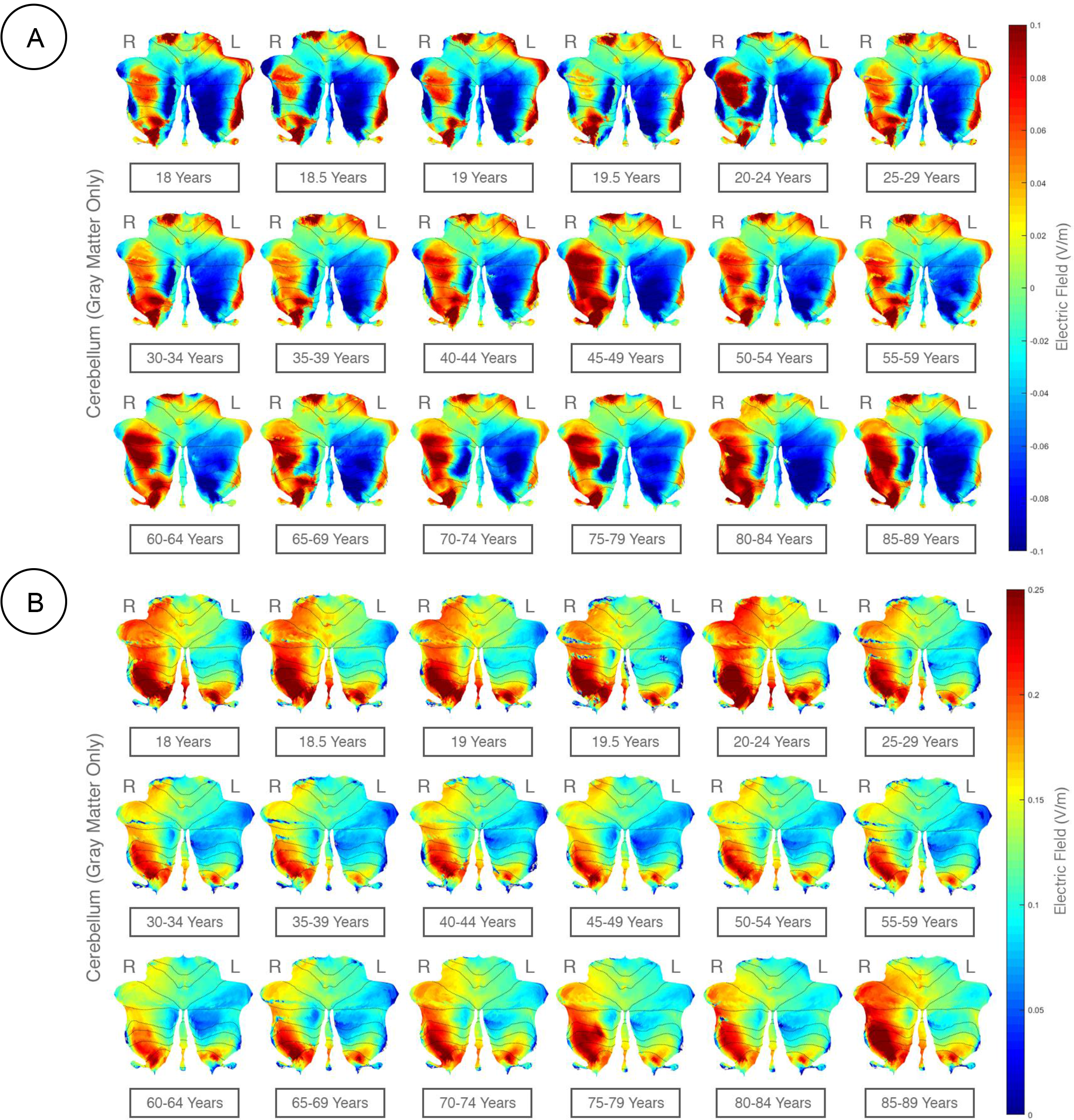
Lobular Electric Field (EF) distribution (as 2D flatmap) in X, Y, Z directions in the cerebellar gray matter. Here, ctDCS using “Celnik Montage” with a direct current of 2mA was simulated to determine the lobular electric field distribution across age-groups. Here, right (R) hemisphere is the targeted side (ipsi) and the left (L) hemisphere is the contralateral side (contra) for ctDCS. Right (R) and Left (L) cerebellar hemispheres are specified in the figure. **A)** flatmap for EF in X direction; **B)** flatmap for EF in Y direction; **C)** flatmap for EF in Z direction.

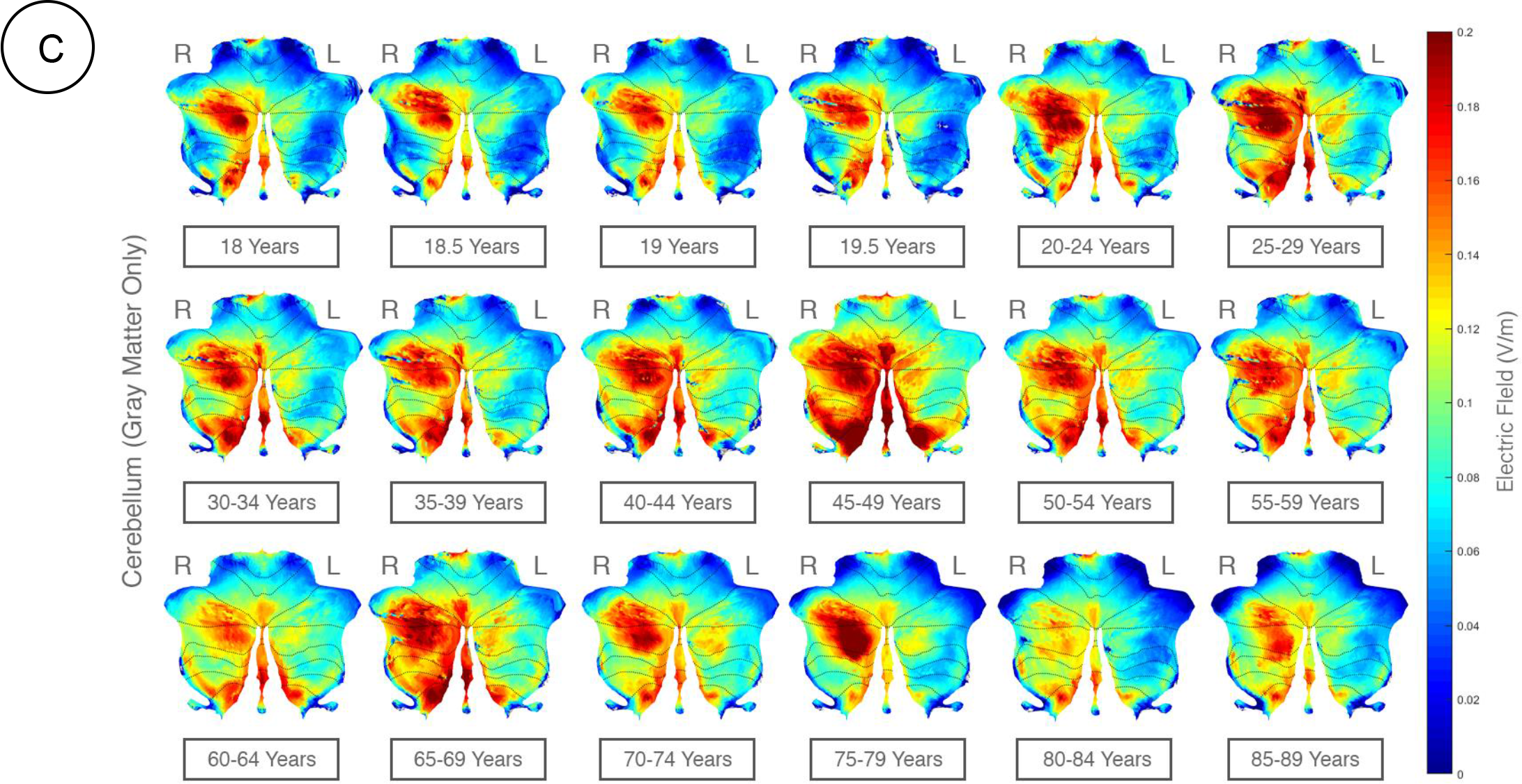

**Appendix 2.**
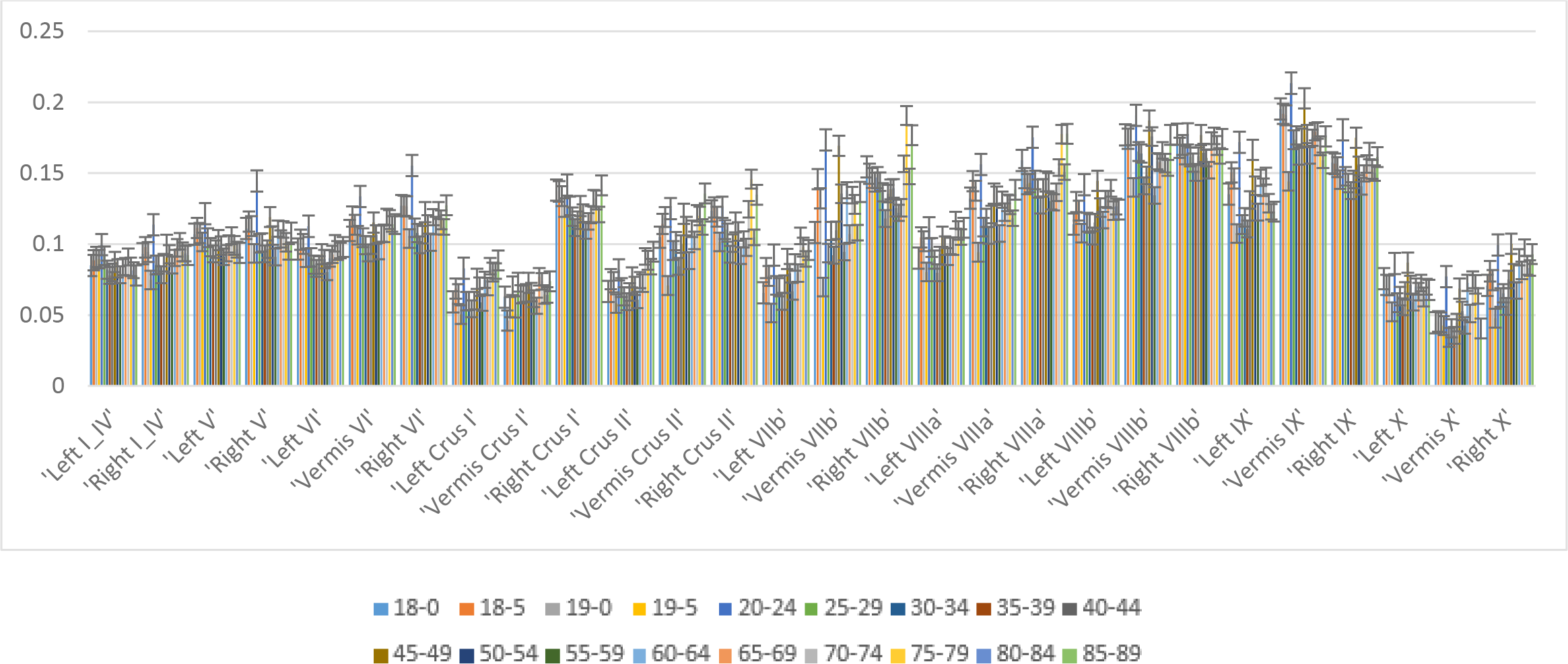
Gray matter electric field strength (V/m) across all the cerebellar lobules.

